# Action imitation via trajectory-based or posture-based planning

**DOI:** 10.1101/2020.09.23.308635

**Authors:** Erica M. Barhorst-Cates, Mitchell W. Isaacs, Laurel J. Buxbaum, Aaron L. Wong

## Abstract

Imitation is a significant daily activity involved in social interaction and motor learning. Imitation has been theorized to be performed in at least two ways. In posture-based imitation, individuals reproduce how the body should look and feel, and are sensitive to the relative positioning of body parts. In trajectory imitation, individuals mimic the spatiotemporal motion path of the end effector. There are clear anecdotal situations in which one might benefit from imitating postures (when learning ballet) or trajectories (when learning to reach around objects). However, whether these are in fact distinct methods of imitation, and if so, whether they may be applied interchangeably to perform the same task, remain unknown. If these are indeed separate mechanisms that rely on different computational and neural resources, a cost should be incurred when switching from using one mechanism to the other within the context of a single task. Therefore observing a processing cost would both provide evidence that these are indeed two distinct mechanisms, and that they may be used interchangeably when trying to imitate the same stimulus. To test this, twenty-five healthy young adults performed a sequential multitasking imitation task. Participants were first instructed to pay attention to the limb postures or the hand path of a video-recorded model, then performed a neutral, congruent, or incongruent intervening motor task. Finally, participants imitated the modeled movement. Spatiotemporal imitation accuracy was greatest after a neutral intervening task, and worst after posture matching. When the primary task involved imitating trajectories, the data suggested a processing cost: movements following the posture-matching intervening task were less consistent with baseline (neutral) performance, suggesting performance may be disrupted by the incongruence. This effect was not observed when imitating limb postures. In summary, we present initial evidence of a partial dissociation between posture matching and trajectory imitation as a result of instructions and intervening tasks that is consistent with the existence of two computationally distinct imitation mechanisms.

## 1. Introduction

Imitation is the process by which an individual reproduces the movements of an observed actor (Heyes, 2021; Thorndike, 1898; Tomasello, 1996; Zentall, 2006). The capacity to intentionally imitate another individual’s actions – such as when copying a movement after some time has passed (Zentall 2006) – is critical to many daily social interactions and is a key component of motor learning and recovery. For instance, learning ballet or the rehabilitation of functional movement after stroke often involve observing and imitating another’s movements. Despite the prevalence of imitation in our day-to-day lives, there is no consensus regarding exactly how people imitate. Much effort to address this question over the last three decades has focused on demonstrating the existence of a mirror neuron network (i.e., neurons that fire for both the observation and performance of the same action; for reviews, see Cattaneo & Rizzolatti, 2009; Molenberghs et al., 2009). However, while the study of mirror neurons has revealed some of the brain regions that may be relevant for imitation, it does not address the question of exactly which features of a motion are being represented during action observation and replicated during imitation. What humans actually reproduce when imitating – and thus, exactly how people imitate – remains unclear.

One prominent hypothesis is that individuals are primarily concerned with copying the configuration of the entire limb in space; that is, matching the postural configurations of one’s own limbs to the limb configurations of the individual-to-be-imitated (Buxbaum et al., 2000, 2014)(Goldenberg, 2009, 2013). For example, when a dancer observes a teacher performing a series of arm movements, she may imitate those movements by copying the positioning of the shoulder, elbow, and wrist at each prominent pose during the movement. We have previously suggested that for imitation of specific configural actions (such as dance postures), people represent movement goals in terms of how their limb postures should look and feel (Howard et al., 2019; Isaacs et al., 2021). Indeed, one well-supported theory proposes that individuals plan all of their movements with the goal of attaining postures that minimize the anticipated discomfort in arm configuration at the end of the movement (Elsinger & Rosenbaum, 2003; Rosenbaum et al., 2001). Much evidence for posture-based imitation stems from research in individuals with limb apraxia, a disorder that commonly results from stroke lesions to the left hemisphere. People with apraxia are frequently impaired in gesture imitation (Buxbaum et al., 2000, 2014)(Goldenberg, 2009, 2013) – especially when the relative positions of body parts are critical (Jax et al., 2006) – despite having otherwise intact motor control. Apraxia is often thought to be related to an inability to form a spatial-configural representation of the body (Sunderland & Sluman, 2000), effects which can be observed when individuals imitate postural configurations with their own body or by positioning the limbs of a mannequin (Goldenberg, 1995). In fact, imitation impairments in these patients have been found to correlate with impairments of the “body schema” (Buxbaum et al., 2000; Schwoebel et al., 2004), a representation supporting the ability to represent the spatial relationships between body parts (Parsons, 1994; Reed & Farah, 1995). This relationship between imitation deficits and impaired body schema representations in patients with apraxia has been cited as evidence that imitation can occur by matching limb postures (Buxbaum et al., 2000, 2014; Goldenberg, 2009, 2013). However, much of this prior work has come from studies that involve imitating static limb positions (e.g., (Goldenberg, 1995)); it therefore remains unclear how well such a mechanism generalizes to more continuous, dynamic movements.

We recently proposed an alternative imitation mechanism that may be particularly well-suited to dynamic movements: copying the motion of the end-effector as it moves through space (Wong et al., 2019). For example, rather than focusing on the configural positioning of her instructor’s limbs at each pose, a ballet dancer might instead copy the flowing motion of the instructor’s hand as it smoothly transitions between poses. Motion along this path forms a “shape” or trajectory in space; we thus refer to this type of imitation as “trajectory imitation.” In this case, the configuration of the individual limb segments to achieve this trajectory are not directly specified as in posture-matching imitation, but are simply those most amenable to producing the desired motion path of the end effector through space; thus trajectory imitation is body-independent (Wong et al., 2015). The ability to imitate by copying trajectories is consistent with the finding that individuals often focus primarily on the end-effector when observing others in preparation for imitation (Matarić & Pomplun, 1998). However, while trajectory imitation has a clear advantage in that the imitator does not need to track the positioning of multiple limb segments over time and may more compactly represent an entire movement as a simple geometric shape, such an approach presents obvious limitations when postural configurations are also relevant to task success. Thus there are clear anecdotal situations in which a person might benefit from imitating postures or trajectories. Whether these are in fact distinct methods of imitation, and whether they may be applied interchangeably to perform the same task, remain unknown, in large part because thus far these two mechanisms have not been directly studied in the context of a single experiment.

Recently, we demonstrated that planning trajectories may enable action imitation, and that some individuals with apraxia are in part impaired at imitating motion trajectories (Wong et al., 2019). In that study, participants with left-hemisphere stroke and neurotypical controls were asked to imitate two-dimensional motion trajectories cued by a human model or via the movement of a cursor (i.e., when no body-posture information was present). We expected that, if people are imitating by copying trajectories, we should see similar imitation performance regardless of whether or not postural information was available. Indeed, the availability of postural information did not affect performance. We interpreted this result as indicating that people could use end-effector trajectory information to guide their imitation efforts, and intriguingly, that when the goal was to imitate trajectories, additional postural information was not beneficial. However, while suggestive, the previous study did not definitively dissociate postural and trajectory based planning. As we did not analyze 3-dimensional arm postures in that task, and as movements were restricted to 2-dimensional space, it is possible that participants simply planned reasonable body postures when body information was not present as long as those postures resulted in hand motion along the displayed trajectory. Indeed, in this and other studies, imitating postures and trajectories will result in the same overt behavior since the postures presented during an action were typically the most comfortable or natural ones to adopt given the end-effector trajectory. As such, the previous study leaves open the question of whether we do in fact employ distinct mechanisms when imitating trajectories or postures, or if people only rely on a single, default mechanism for imitation.

Here, we further test the hypothesis that there are in fact two computationally distinct mechanisms available for imitation. We address the limitations of the prior study by asking participants to perform three-dimensional movements and applied a spatiotemporal kinematic measure to analyze movements of the entire arm rather than just the path of the finger. To probe the question of whether individuals can employ both trajectory- and posture-based imitation, we manipulated the imitation context, influencing which imitation mechanism ought to be used. This context is often intrinsically dictated by the task itself; for instance, when learning classical ballet, a student often begins by focusing on matching the correct arm, torso, and leg positions of the teacher, which results in enhanced posture matching imitation abilities over years of training (Bläsing et al., 2012, 2018; Ramsay & Riddoch, 2001). Based on our prior study (Wong 2019), we reasoned that it should be possible to use instructions to set this context. Indeed, across a variety of motor activities, task instructions have been shown to strongly modulate where individuals focus their attention during a task, consequently affecting task success (for reviews, see (Wulf, 2007; Wulf, 2013) (for a similar effect in stroke patients, see (Kantak et al., 2020)). In these studies, instructions to focus on internal features such as how an individual’s body is moving (e.g., arm postures during a tennis swing) interestingly tend to result in worse performance compared to instructions directing focus toward external features associated with how the environment is affected by the movement (e.g., the motion of the racket and ball). Thus task instructions can have a surprisingly strong influence on the way in which people perform motor tasks. We therefore hypothesized that not only are there two distinct mechanisms for imitation, but that we can influence the choice of mechanism by giving people different imitation goals (i.e., to imitate postures or trajectories), even if the observed movement to be imitated is held constant.

To assess whether participants used both forms of imitation as instructed, we employed a “sequential multitasking paradigm” (Brumby et al., 2018), in which an individual switches between tasks after performing each task for some time, eventually returning to the original task. This task is designed to create a processing cost associated with transitioning from one mechanism to the other, when the intervening task inserted in the midst of a primary task relies on distinct computational processes (Cellier & Eyrolle, 1992; Salvucci et al., 2009). Switching between the primary and intervening task creates large processing demands, and subsequently results in impaired performance when resuming the primary task. Often observed in cognitive tasks, such processing costs can also be measured during motor planning (Orban de Xivry & Lefèvre, 2016). Sequential multitasking paradigms have also been frequently used in action observation studies to assess memory for whether an action has been previously observed or not (for a review, see (Galvez-Pol et al., 2020)); in particular, the sequential multitasking paradigm has been used to demonstrate that the ability to recall static postures and dynamic movements rely on distinct working memory processes (Vicary et al., 2014). To date, however, sequential multitasking paradigms have not been applied to imitation per se.

In the current study, participants first watched a video of a model performing a meaningless upper-arm movement and were instructed that they should either imitate body postures or hand trajectories. Participants then completed an intervening task that was designed to be congruent (no processing cost), incongruent (inducing a processing cost), or neutral with respect to the primary imitation task instructions. We assessed spatial and temporal similarity between the participant and the model as our primary analyses. We also conducted a secondary analysis comparing participant’s movement consistency in the congruent and incongruent conditions against their own performance in the neutral condition. We predicted that in both cases, performing incongruent primary and intervening tasks (e.g., trajectory imitation instructions and the posture-matching intervening task) would induce a larger performance decrement compared to performing congruent primary and intervening tasks (e.g., trajectory imitation instructions and the trajectory intervening task), because of the need in the former case to switch between different processing mechanisms to complete each task. Such findings would provide evidence in support of the existence of two computationally distinct means of imitating – matching postures or copying trajectories – and suggest that individuals are able to flexibly employ either mechanism depending on the task goals.

## 2. Methods

### 2.1 Participants

We recruited participants from Salus University through email solicitation with recruitment procedures approved by the Salus Institutional Review Board. Participants were graduate students or faculty in a healthcare field, such as Occupational Therapy or Optometry. Our sample size was chosen based on previous dual-tasking paradigms that used 20-26 neurotypical participants (e.g., Reed & Farah, 1995). Twenty-five young adults completed the experiment across two sessions (22 female). The average age was 25.0 years (*SD*=4.8, range 22 to 41 years). Participants were right-handed, had full range of motion of the arm, normal or corrected-to-normal vision, and no known neurological disorders. All participants provided written informed consent; experimental procedures were approved by the Einstein Healthcare Network Institutional Review Board. Participants completed two sessions; the first session lasted approximately 1.5 hours, and the second session occurred a minimum of one week later and lasted approximately 1 hour. Participants were compensated $15 per hour for their time.

### 2.2 Materials

For the imitation experiment, participants were seated in a stationary chair 176 cm in front of a 42.5” 4K LG monitor (refresh rate, 60 Hz) mounted on the wall directly in front of the chair, where videos of an actor or static images were displayed using custom experiment scripts written in C++ (https://github.com/CML-lab/DualTaskImitation).

Movement kinematics were recorded using a magnetic motion tracking system (TrakSTAR, Northern Digital Inc., Ontario, CA) with a wide-range transmitter (range, 6 feet) and 4 standard (8mm) sensors. Sensors were placed on the top of the right hand near the wrist, the right elbow joint, and on each shoulder. Movement was recorded at an effective sampling rate of 420 Hz. A piece of tape on the participant’s right thigh, near the knee, served to indicate a home position where the participant should return his or her hand between movements.

Following the imitation experiment, participants were seated at a desk in front of a desktop computer (23.8” Acer R240HY widescreen monitor, 60 Hz refresh rate) and completed a computerized Corsi Block Tapping Task in the Psychology Experiment Building Language (PEBL; http://pebl.sourceforge.net/battery.html).

### 2.3 Procedure

The imitation experiment consisted of a sequential multitasking paradigm (Brumby et al., 2018; Cellier & Eyrolle, 1992; Salvucci et al., 2009) in which participants were asked to watch videos of an actor generating a movement with what appeared to be his left arm, complete an intervening task, then imitate the movement they had watched using their right arm (i.e., imitating as if the movement was observed in a mirror, which is considered a more natural form of imitation; (Bekkering et al., 2000; Franz et al., 2007); see Figure 1). Movements of the actor were in fact recorded using the actor’s right arm to ensure greater comparability to the participants’ own movements; videos were then mirror-reversed to make it appear as if the actor was using his left arm. For the imitation portion of the task, movements consisted of two or more static postures (with a ~1 second hold at each posture) with dynamic transitions between those postures, and were designed to have no obvious meaning associated with the movement. For example, one movement would consist of lifting the arm off the lap and bending it into a raised “L” shape out to the side (posture 1), a brief pause, extending the arm out to the side to a low diagonal (transition to posture 2), a brief pause, and then returning the arm to the lap (see Figure 2). Postures and transitions involved motion about the shoulder and elbow, but did not include flexion/extension at the wrist or movement of the fingers; the hand remained in a fist throughout the motion. Movements included 1 or 2 dynamic transitions (2 or 3 static postures, respectively), with the transition movements being either straight or curved. After watching the movement and completing an intervening task (see below), participants were instructed to wait until the go cue, in which the word “Imitate!”was displayed on the screen along with an auditory tone, to begin imitating. The go cue appeared 500 ms after the end of the intervening task. If participants moved before the go cue (detected as movement of the hand more than 5 cm from the home position), the trial restarted.

**Figure 1.**
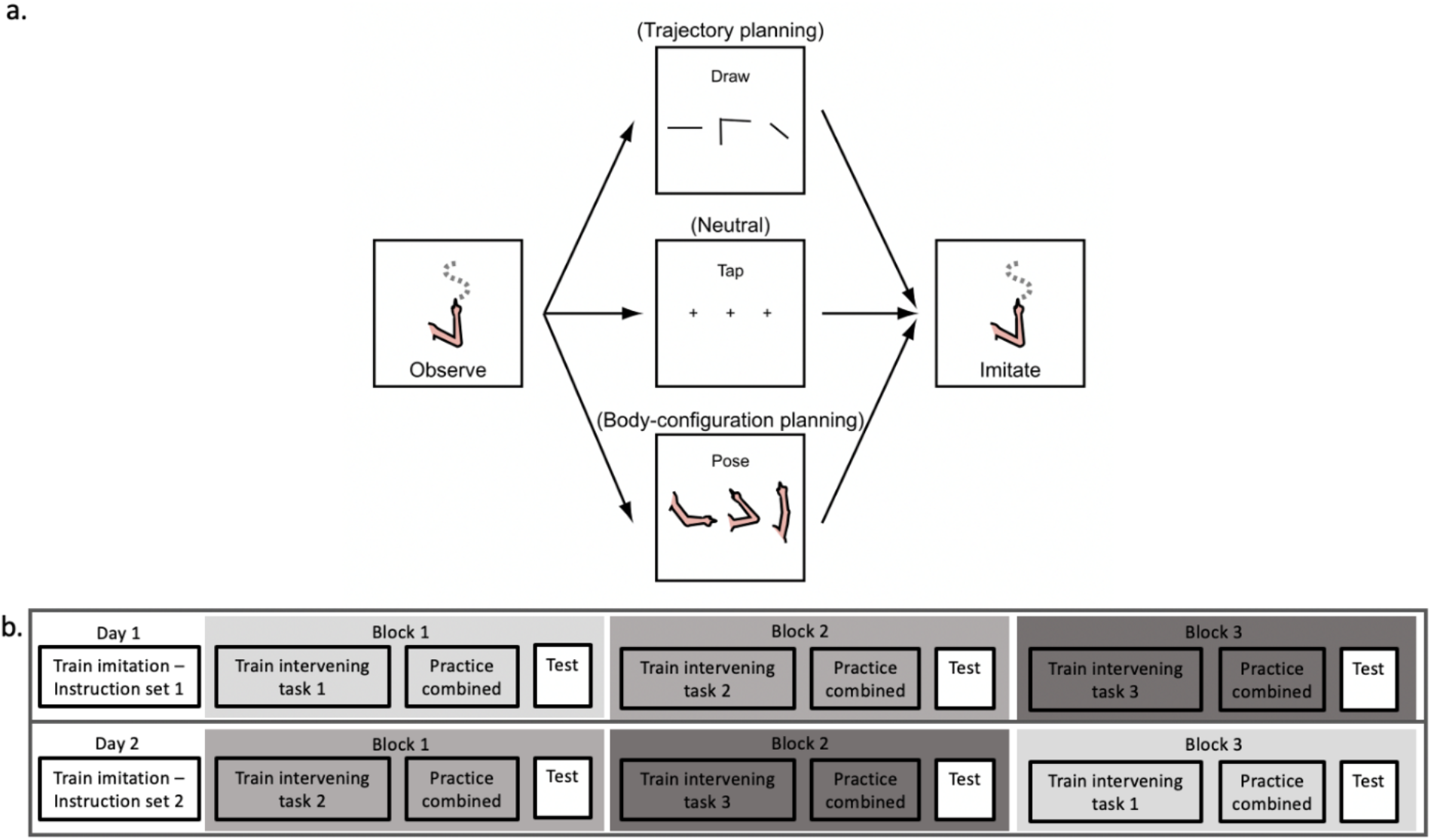
Overview of experimental paradigm. Panel a. overviews the sequence of events for each trial: first, observation of the primary task movement, second, completion of the intervening task, third, performance of the primary task movement. Panel b. overviews the sequence of events across each testing session. Each participant completed all three intervening task conditions (represented as the shading of the blocks; e.g., dark gray could represent the neutral IT condition) on both day 1 and day 2 of testing in a counterbalanced order. <u>On Day 1 and Day 2, participants received different instructions about what movement feature (postures or trajectories) to focus on when imitating</u>.

**Figure 2.**
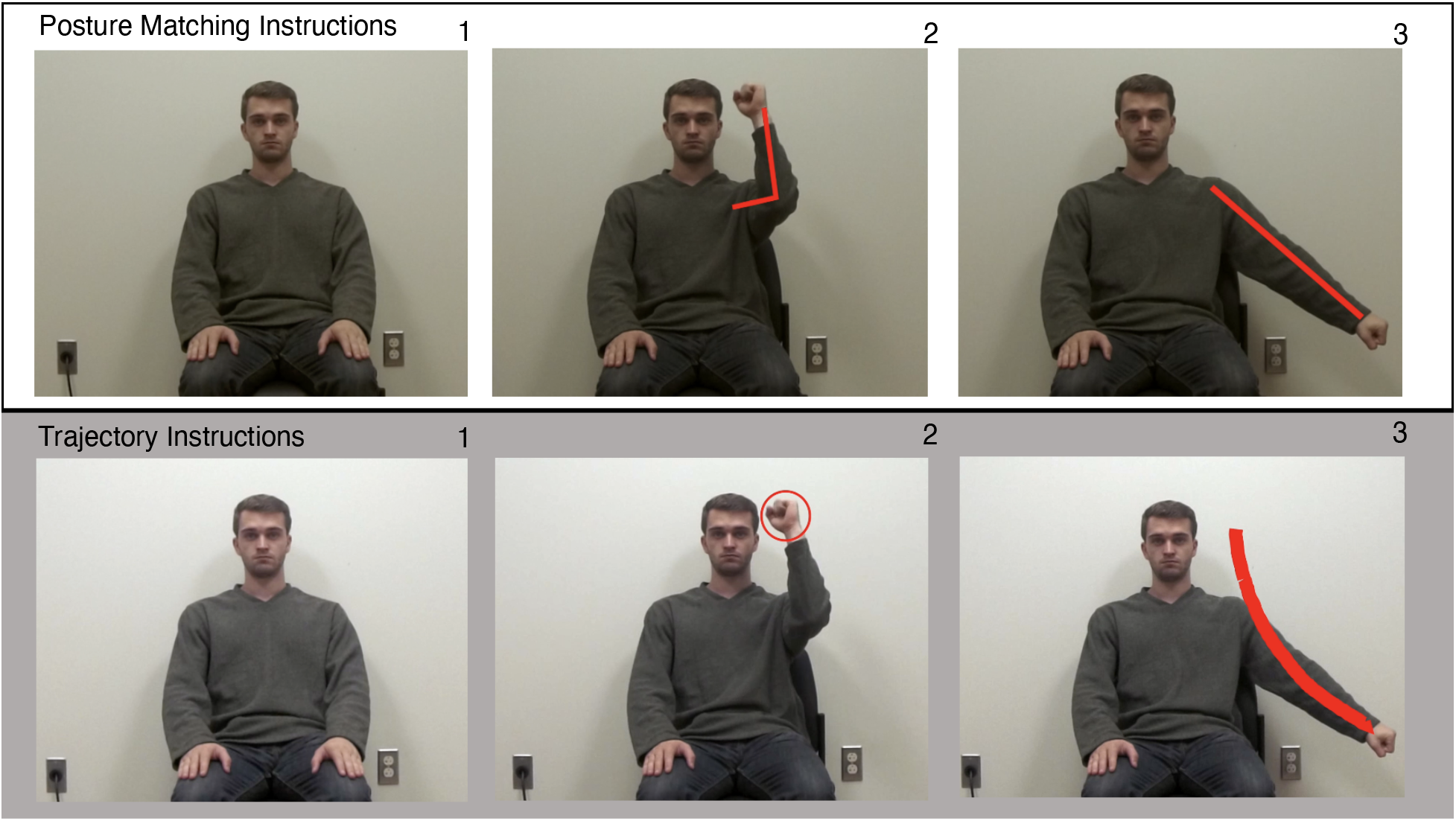
Snapshots of Instruction videos. The top panel is an example of the instruction video provided in the Posture Matching PT. Participants watched the model perform a two-posture movement, pausing at each position. Emphasis lines appeared to draw attention to the position of the body parts (wrist, elbow, shoulder). The bottom panel is an example of the instruction video provided in the Trajectory PT. Participants watched the model perform a two segmented movement with the circle tracking the movement of the hand (bottom left and bottom middle). Upon reaching the end position, a “trace” of the movement appeared, which represented the path the hand took through space (bottom right). Emphasis lines were only present during the initial four practices on each day of testing. Photos of author M. Isaacs used with the author’s permission.

On each day (Fig. 1b), participants were given a specific set of instructions for the entire session about what movement features to focus on during imitation (see Instructions manipulation below). They then completed 4 imitation-only practice trials in which they watched a video of the model performing the movement, and were immediately instructed to imitate the observed movement with no intervening task present. Next, participants completed 3 blocks in each session, one for each of the 3 intervening tasks described below. Each block began with 10 familiarization trials of the intervening task to be performed in that block. This was followed by 4 combined practice trials, where they first watched the model movement without moving their hand, completed the same intervening task they had just practiced, then performed their imitation of the model movement upon seeing a go cue. Finally, participants completed 20 test trials (analogous to the combined practice trials) to finish the block, and received a brief (~30 seconds) break before beginning the next block. Following the completion of all blocks in the session, participants completed the Corsi task.

#### 2.3.1 Primary task: Instruction manipulation

The primary task (PT) was to imitate gestures. Instructions were used to manipulate the focus of attention during imitation on either limb postures or end-effector trajectories. That is, participants received a different set of instructions in each session (order counterbalanced between subjects). For each instruction, the practice videos were edited to emphasize the relevant set of instructions. In the Posture Matching instructions (Posture Matching PT), the experimenter instructed participants to: “Pay attention to the position of the body parts—for example, how the elbow, shoulder, and wrist are positioned.” In the practice videos, when the model reached each static posture, the movement was momentarily paused and straight red lines appeared on the model’s forearm and upper arm to encourage participants to pay attention to the positions of the body parts relative to each other (Fig. 2, top panel). In the Trajectory instructions (Trajectory PT), the experimenter instructed participants to “Pay attention to the path that the hand takes through space.” In the practice videos, the model’s hand was encircled by a red circle that followed the movement of the hand; upon completion of the movement, the full path of the movement was displayed as a red line on the screen to encourage participants to attend to the path of the end effector (Fig. 2, bottom panel). These explicit instruction cues <u>(i.e., the red lines in the videos)</u> were not present during test trials. The experimenter also verbally reminded participants to either “focus on the position of the body parts” or to “pay attention to the path the hand takes through space” at 7 points during each session: prior to the instruction-only practice videos at the very beginning of the session, prior to the combined practice trials (one for each of the three intervening tasks described below), and prior to the test trials for each intervening task. In all cases, participants were instructed to imitate the movement at the same speed as the model.

In order to minimize learning effects, we randomized the order of the movements within easy (the first 8 trials, consisting of 2 static postures each) and hard (the second 12 trials, consisting of 3 static postures each) trials between blocks so that participants did not view the movements in the same order each time. We also created forward and reversed videos of each movement, which contained equivalent kinematic information (i.e., postures and trajectory shapes) but allowed us to minimize the number of repeated exposures to the exact same movements. Reversed videos were generated by running time backwards in the videos. Participants saw either the forward or the reversed videos in the first session (paired with either Posture Matching or Trajectory PTs), and then the other set of videos in the second session, paired with the other set of instructions. We hypothesized that the direction of movement would not significantly influence imitation accuracy. For trials where the video was reversed, we also inverted time in the kinematic recordings from the participants such that in our analyses the recorded gestures from all trials and all participants were compared to the kinematics of the forward-in-time model movement. Note that the model kinematics were the exact kinematics recorded from the model during filming of the movement videos for this experiment, using the same trakSTAR and placement of trackers.

#### 2.3.2 Intervening tasks

We included three intervening tasks (IT), which were manipulated within-subjects. The order of ITs was randomized and counterbalanced between participants and between sessions. These tasks were meant to evoke processing interference with the primary imitation task; as such one task was related to planning limb postures and a second was related to planning trajectories. The third task served as a control for the working memory demands associated with remembering the PT while performing an IT. In these three ITs, participants were required to produce a movement once every 1200 ms in time with a metronome for 6 iterations before returning their finger to the home position.

In the Posture Matching IT, participants were asked to reproduce three body postures presented in pictures on the screen, from left to right, in time with the metronome (see Figure 3). In the Trajectory IT, participants saw three meaningless shapes on the screen, and were asked to extend their right arm directly in front of them and draw the shapes in the air using their whole arm in time with the metronome. Examples of these shapes include a straight horizontal line, a rotated “L”, and a diagonal line. Drawing movements were to begin at the dot in each image (see Figure 3).

**Figure 3.**
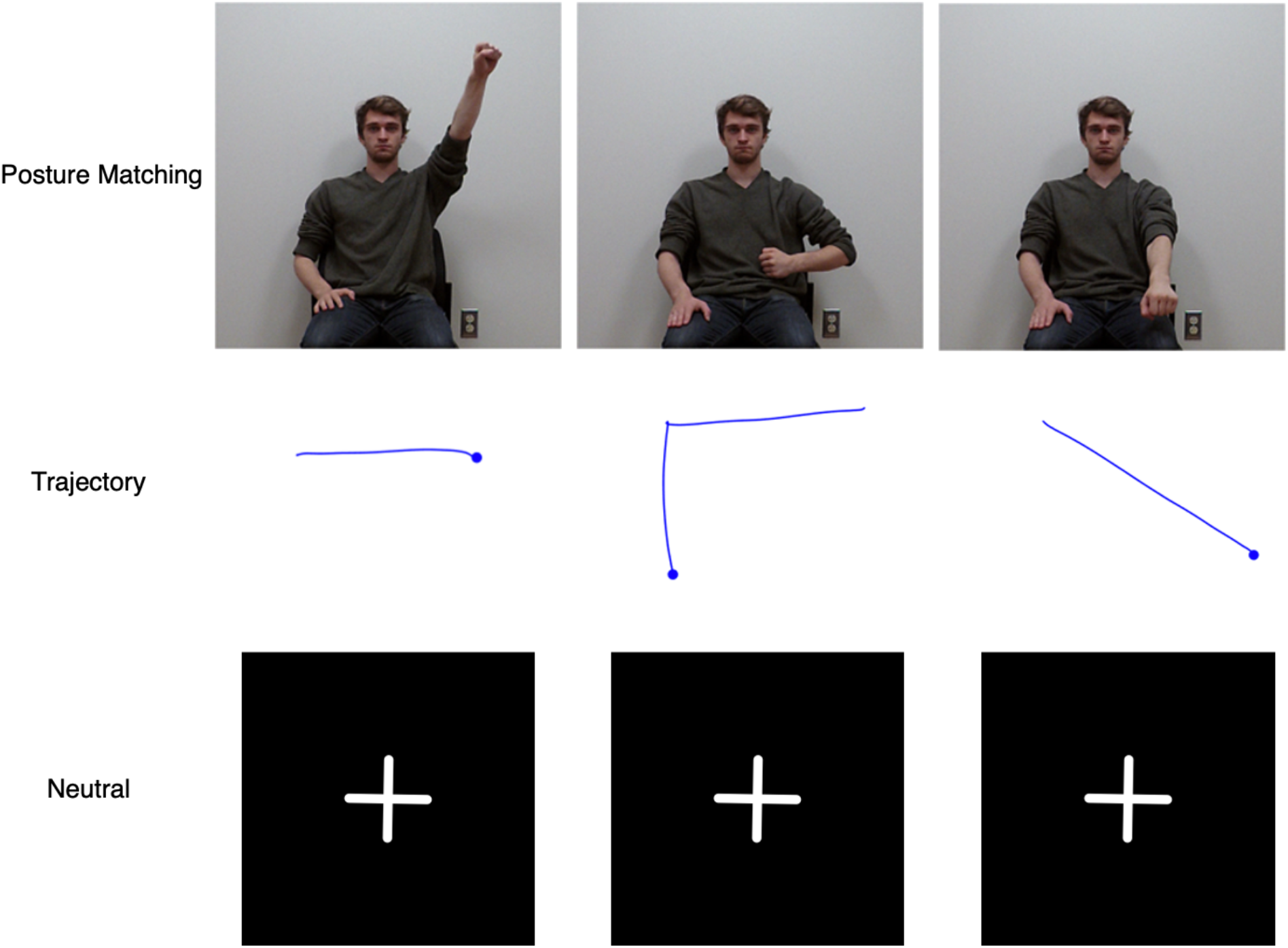
Intervening Tasks. Participants completed each of three ITs for both sets of Instructions. In the Posture Matching IT (top row), participants matched the arm positions in the photographs of the model from left to right in time with a metronome that presented a tone once every 1200 ms. They heard 6 tones and were asked to match the postures from left to right twice through. Posture Matching IT is congruent with Posture Matching PT, but incongruent from Trajectory PT. In the Trajectory IT (middle row), participants raised their hand in front of them into the air and drew the shapes one at a time from left to right in time with the metronome. They completed the row of shapes twice for 6 metronome tones. We instructed participants to start the drawing movement at the circle on each shape. The Trajectory IT is congruent with Trajectory PT, but incongruent from Posture Matching PT. In the Neutral IT (bottom row), participants tapped their hand on their upper leg in time with the metronome, 6 times. We expected the Neutral IT to be unrelated to either Posture Matching or Trajectory PT and to serve as a control for working memory demands. Photos of author M. Isaacs used with the author’s permission.

To control for working memory demands associated with the delay that the intervening tasks created between watching the action and performing the imitation, we included a neutral intervening task in which participants saw 3 fixation crosses presented side by side on the screen and simply had to tap their hand on their thigh in time with the metronome – a motor response that does not preferentially invoke the planning of either body configurations or trajectories. Fixation crosses were used to provide an analogous amount of visual cuing information across conditions. When tapping, participants were instructed to slide their hand up their leg away from the home position, and keep the heel of their hand on their leg.

#### 2.3.3 Corsi task

As our task required individuals to hold a multi-segment movement in mind while performing an IT, we wanted to control for potential individual differences in spatial working memory abilities. To assess spatial working memory, participants completed a standard computerized Corsi block tapping task at the end of each session. Participants viewed a layout of nine blue squares on a black background. Blocks briefly turned yellow one at a time in sequence, and after the last block in the sequence returned to blue, participants were instructed to reproduce the sequence by clicking on the boxes in the same order. Participants first completed 3 practice trials, then began the experimental trials. Sequences progressed in difficulty starting with 3 blocks. If participants correctly reproduced at least one of the two sequences at that length, the sequence became longer. When participants failed both attempts, the task ended. The maximum number of blocks correctly reproduced was used as that individual’s Corsi block span. For each individual, we took the average of their two Corsi span assessments. The average Corsi span across individuals was 6.68 (*SD*=1.11, range 5 to 9).

### 2.4 Data Analysis

Kinematic data were analyzed in Matlab using custom scripts. Movement start and end were automatically identified according to a velocity threshold of 5 cm/s. These points were verified by visual inspection and manually adjusted if necessary by one researcher, then they were independently confirmed by a second researcher. The start of each movement was identified as the point at which participants initiated a movement away from the first static posture, and the end of each movement was the point at which participants arrived at the final static posture; that is, we only included the “core” portion of the movement from the first to the last target postures, excluding motion to and from the home position. As a preliminary analysis, we counted the number of movement segments executed on each trial using the same velocity threshold method to mark the movement segments. Segment-count errors were identified when there were more than or fewer than the correct number of movement segments generated during that trial. We observed a greater likelihood of producing movement segment errors after the Posture Matching or Trajectory ITs compared to the Neutral IT, suggesting that the Neutral IT was indeed “neutral” with respect to the primary imitation task^1^.

#### 2.4.1 Primary task performance

Accuracy in the primary task (imitation performance) was measured by quantifying the degree of dissimilarity between the participant’s kinematics and the model’s kinematics (analysis 1). We also compared the participant’s kinematics in one condition to their own kinematics for the same movement in another condition (analysis 2) to examine whether the intervening task disrupted individual participant’s performance relative to their imitation ability in the neutral condition. For these analyses we used a Procrustes Distance (PD) metric, which is a measure of dissimilarity between two geometric shapes after accounting for any affine transformations (i.e., translation, rotation, and scaling) (Goodall, 1991). A larger PD indicates greater dissimilarity, or error, compared to the referent.

To describe the motion of the entire arm throughout the movement (including the path of the hand), we projected all data into a coordinate system defined by the orientation of the participant’s body in space (where x points to the participant’s right, y points forward, and z points up). We then developed a novel method of summarizing the configuration of the arm by treating the arm as a plane rotating in 3D space, with its origin at the shoulder, y axis directed along the shoulder-elbow vector, z axis directed along the normal to the plane, and containing the elbow-hand vector (Figure 4). We calculated the instantaneous orientation of the plane as a quaternion relative to the body-centered coordinate frame. Two additional angles were needed to define arm orientation fully; these consisted of the elbow angle (the angle from the shoulder-elbow to the elbow-hand vector), and the roll angle of the hand within the plane (reflecting pronation/supination of the forearm, defined as the angle between the normal to the hand sensor and the normal to the plane). These latter two angles were normalized by 360 degrees to put them on a comparable scale to the arm-plane quaternion. This six-element vector was computed at every time step, and the entire high-dimensional trajectory was then time-normalized to have 200 equally spaced points. Finally, we compared this time-normalized arm representation against a similarly constructed one for the model. For this comparison we used a modification of the PD algorithm in which movements could be scaled differently in each dimension, as it improved the quality of the fits (Rohlf & Slice, 1990).

**Figure 4.**
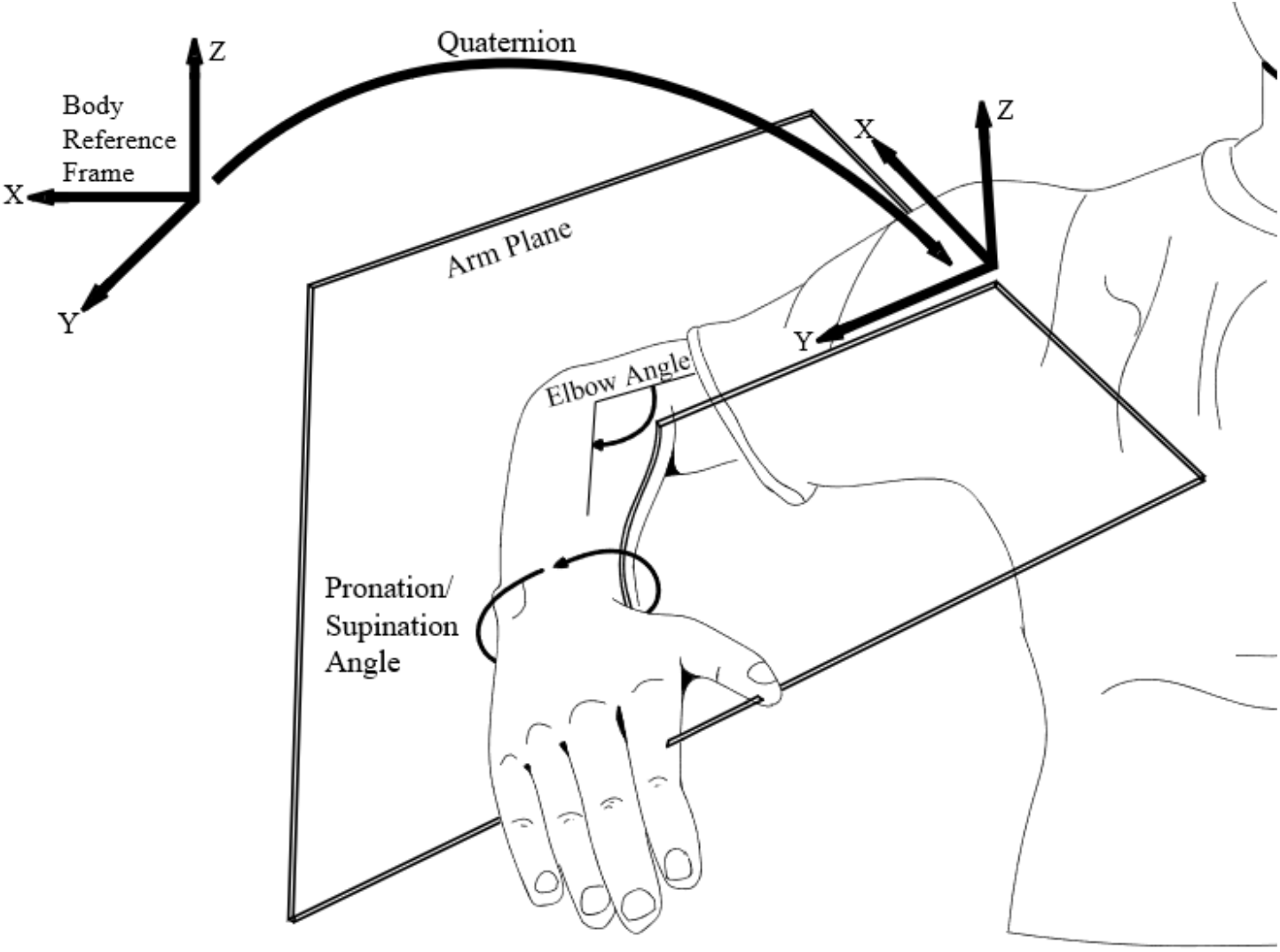
Arm configuration at each time step was described using six values: a quaternion that described the rotation of the axes of an “arm plane” (defined as the plane containing the shoulder-elbow and elbow-wrist vectors) relative to a body-centered reference frame (4 values); the angle of the elbow within the arm plane (1 value); and the pronation/supination angle of the forearm (1 value). The latter two angles were normalized by 360 degrees to be of a similar scale as the individual elements of the quaternion.

PD measures are typically bounded between 0 and 1, where 1 reflects maximal dissimilarity (Goodall, 1991; Gower, 1975). However, when the algorithm is modified to allow each dimension to scale differently as suggested by Rohlf and Slice (Rohlf & Slice, 1990), PD values may exceed 1. Nevertheless, even in such cases PD values remain relatively constrained to the range 0-1 such that PD values greater than 1 reflect unusually large behavioral outliers; we therefore removed these outliers in our analyses. This resulted in the removal of 0.03% of our data (8 trials) for analysis 1 and 0.3% of the data for analysis 2 (69 trials).

#### 2.4.2 Movement time

As participants were instructed to imitate movement speed as well as spatial position, we also assessed temporal accuracy errors (analysis 1b). Timing errors were computed as the absolute difference between the time taken for the participant and the model to produce a given movement. Larger values indicate a greater difference between participant and model movement times, suggesting more timing dissimilarity. The average model movement time for all movements was 2.93s with a range from 1.36 to 4.60s.

#### 2.4.3 Statistical analysis

All statistical analyses were performed in R (R Core Team, 2013) We used the *lme4* package (Bates et al., 2015) to fit generalized linear mixed effects models, with significant effects of interest identified by calculating likelihood ratio tests between models with and without the factor of interest. Because the PD data are bounded and non-normally distributed, we used a generalized linear model with a log link to analyze imitation accuracy. For the Movement Time data, we used a square root link, which normalized the data to resemble a more normal distribution. Because of experimenter or technical error, 86 of 3000 trials (2.87%) were missing. Our hypothesized models included fixed effects of Instruction (2 levels), Intervening Task (3 levels), and the Instruction*Intervening Task interaction (2×3), along with a random effect of Participant. We also checked for an effect of instruction Order and video Direction (forward or reverse) in our models; we hypothesized *a priori* that neither of these factors would have a significant impact on our data. Finally, we tested whether performance was influenced by spatial working memory span. For all statistical models, we performed pairwise contrasts with the *emmeans* package (Lenth, 2019) using a Tukey adjustment for multiple comparisons.

### 3. Results

#### 3.1 Imitation is disrupted by an imitation-related intervening task

In our first analysis we compared participants’ imitation to the model to assess the extent of spatial dissimilarity using the Procrustes Distance algorithm. We first checked if there were any effects of video Direction and instruction Order. As hypothesized there was no significant effect of Direction (χ^2^(1)=0.38, *p*=0.5) or Order (χ^2^(1)=0.38, *p*=0.5), so we removed these terms from the model. The resulting model suggested a trending effect of Intervening Task (χ^2^(4)=8.60, *p*=0.07) but no effect of Instruction (χ^2^(3)=0.97, *p*=0.8). There was also no significant interaction (χ^2^(2)=0.92, *p*=0.6). While not significant, we noted that imitation following the Neutral IT tended to be best (lowest PD) overall (see Table 1 for means in each condition). Adding average Corsi block span to the model did not improve model fit (χ^2^(1)=1.29, *p*=0.3), suggesting that imitation accuracy was not significantly related to spatial working memory.

**Table 1.** Average imitation performance compared to the model.

| Intervening Task | Posture Matching Primary Task |  |  | Trajectory Primary Task |  |  |
| --- | --- | --- | --- | --- | --- | --- |
|  | Posture Matching | Trajectory | Neutral | Posture Matching | Trajectory | Neutral |
|  | *M (SD)* | *M (SD)* | *M (SD)* | *M (SD)* | *M (SD)* | *M (SD)* |
| PD | .111 (.137) | .103 (.119) | .099 (.122) | .114 (.145) | .108 (.130) | .096 (.128) |
| Movement Time (s) (unsigned) | .831 (.623) | .777 (.608) | .724 (.594) | .844 (.625) | .785 (.614) | .696 (.564) |
| Movement Time (s) (signed) | 0.742 (.728) | 0.595 (.792) | 0.590 (.727) | 0.744 (.750) | 0.688 (.726) | 0.597 (.674) |

Note. Movement Time represents the difference in the time taken for the model and the participant to perform the movement.

Participants were also instructed to imitate movement timing (movement speeds and static posture hold times) as well as position; thus we additionally examined how well participants were able to imitate total movement time by comparing the absolute difference between the time taken for the participant and the model to produce a given movement^2^. Larger values indicate a greater timing dissimilarity (i.e., more error) between participant and model movement times. We first checked if there were any effects of video Direction and instruction Order. There was a significant effect of Direction (χ^2^(1)=50.29, *p*<0.0001) but not of Order (χ^2^(1)=0.85, *p*=0.4). We therefore retained Direction as a nuisance variable in our analysis. Using this model, we observed a significant effect of Intervening Task (χ^2^(4)=25.57, *p*<0.001), but no significant effects of Instruction (χ^2^(3)=0.82, *p*=0.8), nor was there an Instruction*Intervening Task interaction (χ^2^(2)=0.81, *p*=0.7). Planned pairwise contrasts revealed that imitation movement time errors following the Neutral IT were significantly smaller than Movement Time errors following the Posture Matching (*t*=-7.29, *p*<0.0001) and Trajectory ITs (*t*=-4.16, *p*=0.0001). Movement time errors following the Posture Matching IT were significantly larger than following the Trajectory IT (*t*=3.13, *p*=0.005). See Table 1 for means. These data reveal that individuals were more accurate at imitating movement timing following the Neutral IT than the two interference conditions, and moreover that imitating movement time was more accurate after Trajectory than Posture Matching ITs. Taken together with measures of spatial imitation accuracy, these results suggest that in general, having to perform an intervening task related to either of the hypothesized mechanisms supporting imitation is disruptive to the ability to imitate both spatial and temporal movement features.

### 3.2 Within-subjects consistency reveals a processing cost for imitating trajectories

In the prior analysis, we assessed how accurately people imitated in contrast to the ideal (model) movement. While we did not observe the expected interaction between congruent versus incongruent primary and intervening tasks, this may have been due to a large amount of variance unrelated to our manipulations. That is, because it may be challenging in general to exactly reproduce the kinematics of an observed model, our measure of accuracy compared to the model may not have been sufficiently sensitive to detect any additional behavioral variance arising from the IT manipulation. Thus as an additional, potentially more sensitive measure of imitation performance, we also examined the ability for any given individual to repeatedly produce the same movement consistently in all conditions. In other words, the disruption induced by an intervening task might be better detected by comparing each participant’s performance for a given movement shape and condition to their own performance of the same movement shape in a different condition (i.e., a measure of variance versus a measure of bias). Specifically, since the same set of movements were imitated in every block, and because in our prior analyses we confirmed as predicted that the Neutral condition is least disruptive of imitation accuracy, we calculated a PD value that represented a participant’s performance dissimilarity for imitation following the Posture Matching and Neutral ITs, and following the Trajectory and Neutral ITs, within each Primary Task instruction condition. A higher PD value indicates less imitation consistency with reference to the neutral condition, suggestive of greater disruption due to the IT.

We again ran a generalized linear mixed effects model with fixed effects of Instruction (Posture Matching or Trajectory) and Intervening Task Comparison (Posture Matching vs. Neutral or Trajectory vs. Neutral) as well as the Instruction X Intervening Task Comparison interaction. There was no significant effect of Order (χ^2^(1)=0.98, *p*=0.3) but there was a significant effect of video Direction (χ^2^(1)=5.33, *p*=0.02). We dropped Order and retained Direction as a nuisance variable in the model. With this model, there was no significant effect of Instruction (χ^2^(2)=4.95, *p*=0.08), and no significant effect of Intervening Task (χ^2^(2)=4.94, *p*=0.08), but there was a significant interaction (χ^2^(1)=4.52, *p*=0.03). Planned contrasts revealed a significant difference between Intervening Task comparison types for the Trajectory PT (*z*=1.97, *p*=0.049) but not for the Posture Matching PT (*z*=-1.01, *p*=0.3). As depicted in Figure 5, for the Trajectory PT, there was greater disruption in imitation behavior following the Posture Matching IT than following the Trajectory IT. In other words, the combination of Trajectory PT and Trajectory IT resulted in performance that was relatively less disrupted (i.e., more consistent with performance following the Neutral IT) compared to combination of Trajectory PT and Posture Matching IT. This effect was present regardless of the spatial working memory span of individual participants: adding Corsi block span to the model did not improve model fit (χ^2^(1)=0.41, *p*=0.5). These results are consistent with our hypothesis that participants exhibit greater imitation performance decrements (i.e., larger processing costs) when the instructions and intervening task are incongruent, particularly when imitating trajectories.

**Figure 5.**
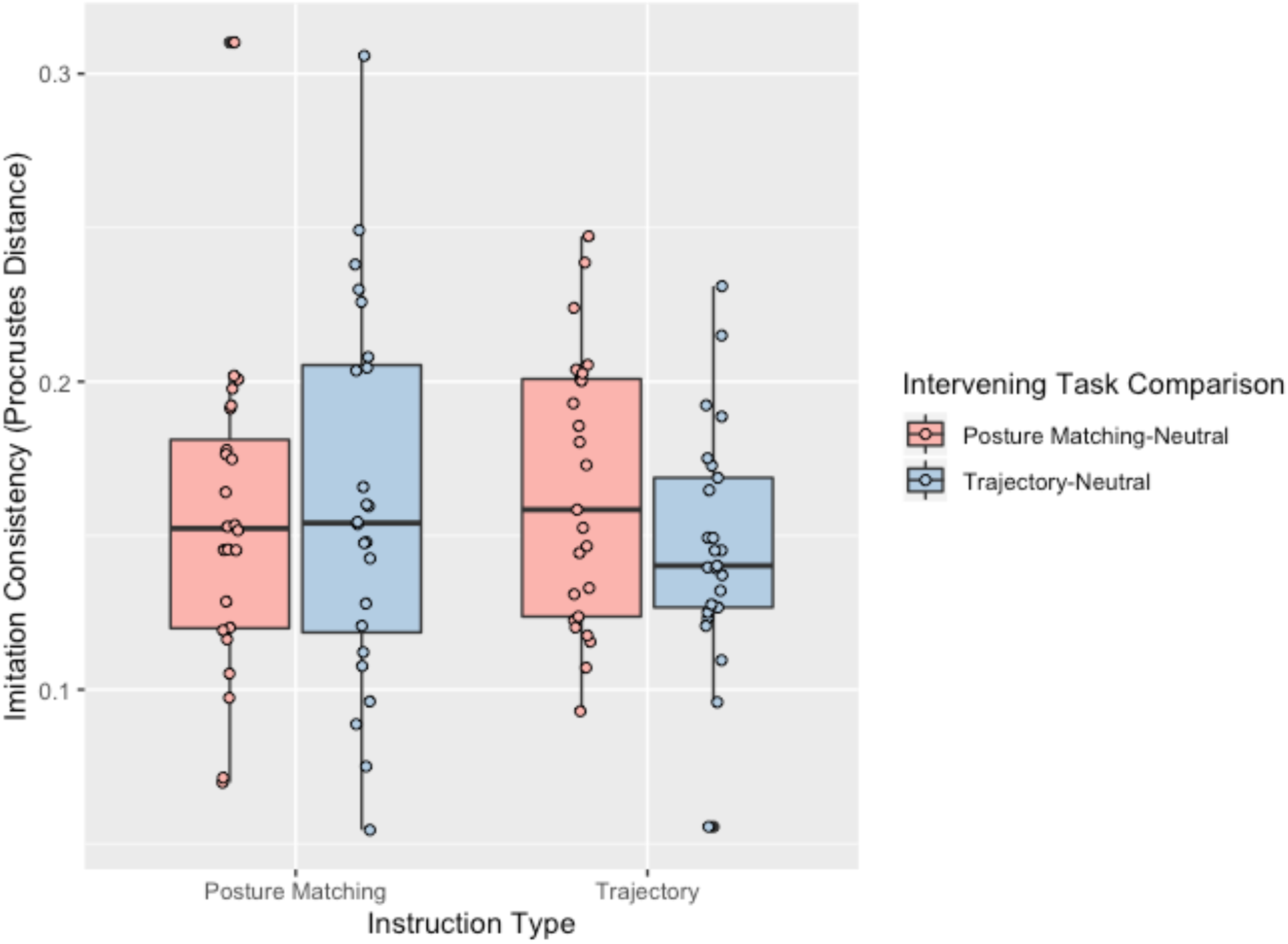
Boxplot of Procrustes Distance results for imitation consistency. A larger number indicates more dissimilarity within an individual comparing imitation following one of the interfering intervening tasks to the Neutral IT.

## 4. Discussion

In the current study we aimed to test whether individuals could imitate the same stimuli via either posture matching or trajectory mechanisms using a sequential multitasking paradigm. We reasoned that if people can imitate using both mechanisms, we should observe a cost when switching from one mechanism to the other, such as when performing an intervening task that requires a different processing mechanism compared to the primary task. Our primary task was to imitate gestures by attending to either the limb postures or path of the hand of the model. In tandem, participants also completed one of three motor intervening tasks: two that were congruent/incongruent with the instructions and designed to potentially create interference, and one that served as a neutral baseline. In our first analysis (examining imitation accuracy in comparison to the model), we saw no overall effect of the primary task, but did observe that the congruent and incongruent intervening tasks tended to be more disruptive to imitation as suggested by worse spatiotemporal imitation accuracy relative to the neutral intervening task. When we then controlled for individual subjects’ variability by measuring within-subjects consistency (i.e., examining how much the IT disrupted imitation compared to the best performance achievable following the neutral IT), we were able to observe a processing cost in the trajectory primary task condition. Specifically, when participants had to complete a posture-matching intervening task that was incongruent with the primary trajectory imitation task, their performance was less consistent with their own baseline performance. These effects were unrelated to any potential individual differences in spatial working memory as assessed by Corsi span (for similar lack of working memory effect, see (Schütz & Schack, 2020)). Each of these results is discussed below in turn. Together these results present initial evidence that switching between trajectory and posture matching imitation tasks may be computationally burdensome and induces a processing cost, in line with our hypothesis that people can imitate using either of these two distinct methods depending on the task context.

In our first analysis of spatiotemporal accuracy compared to the model, our results were only partially consistent with our hypothesis. While we expected an interaction between primary and intervening tasks that would provide evidence of a processing cost, we only observed an effect of intervening task. The posture matching and trajectory intervening tasks were intended to be cognitively and motorically disruptive of the primary task, whereas the neutral intervening task was intentionally designed to obviate these spatio-motor demands, predominantly preserving only the delay between observation and imitation. Our results suggest that indeed, performance of the primary imitation task following the neutral intervening task was most similar to the model, regardless of the task instructions. In contrast, the posture matching and trajectory intervening tasks both elicited a “cost” compared to the neutral intervening task in the form of decreased spatiotemporal accuracy, suggesting that these intervening tasks disrupted imitation. Additionally, we noted that imitation following the trajectory intervening task was more temporally accurate than imitation following the posture matching intervening task, although we did not observe an analogous pattern for spatial accuracy. Overall, this suggests that the neutral intervening task was the least disruptive to imitation, while the posture-matching intervening task was most disruptive.

In our second analysis we then compared participant’s performance following each intervening task to their own baseline (neutral) performance. Unlike in our measures of imitation accuracy, this measure of consistency yielded clear evidence of a processing cost when the primary task was to imitate trajectories (we discuss why we may not have seen an analogous processing cost for imitating postures below). Switching to a posture matching intervening task elicited a “cost” that impaired performance of the primary task. We suspect that the reason this effect may not have been observed in our original accuracy measures is because we were actually measuring two kinds of errors in that case: errors of imitation more generally, and errors related to potential disruptions by the intervening task. The former source of error could arise from needing to attend to and perform a mirror mapping of the model’s movement to one’s own body, maintain the components of the movement in working memory, and execute the movement – each of which would introduce “noise” that would be relatively independent of the specific imitation condition but would be relatively constant across conditions for a given individual. By comparing each person’s performance to their own “baseline” performance (i.e., by assessing movement consistency), we could account for these sources of error and instead more directly examine specific disruptions arising from the intervening task. Thus by adopting a within-subjects analysis, we were better able to detect the presence of a processing cost when participants had to switch between trajectory imitation and a posture matching intervening task (Cellier & Eyrolle, 1992).

Our findings are consistent with prior research that has posited the existence of at least two distinct ways to imitate, with evidence for both posture matching (Buxbaum et al., 2014; Goldenberg, 2009, 2013) and trajectory matching (Wong et al., 2019) mechanisms. These two dissociated processes are hypothesized to emphasize different aspects of the movement-to-be-imitated: posture matching requires the ability to represent how the body should look and feel during and at the end of the movement, whereas trajectory matching instead requires the ability to represent end-effector motion in a body-independent manner. As these are both useful but distinct means of imitating movements, we hypothesized that people may be able to make use of both mechanisms. Our current study is the first to our knowledge to directly test for the existence of these two mechanisms in the same paradigm (and using the same stimuli), to determine whether people can indeed imitate in both ways depending on task demands.

In our daily lives, there are clearly conditions that favor the use of one mechanism over the other. For example, body postures and joint angles are critical to imitation success when learning ballet or modern dance (Bläsing et al., 2012, 2018; Ramsay & Riddoch, 2001), or in the static posture imitation tasks that are often used with clinical neurological populations to test for apraxia (e.g., (Goldenberg, 1995, 2009, 2013)). In contrast, trajectory imitation tasks may be more favorable when one needs to copy a path, such as drawing the solution to a maze or navigating around obstacles. Some previous research has even suggested that an observed action may be represented in terms of both postural forms and motion trajectories (Giese & Poggio, 2003).

We note, however, that a trajectory-matching method of imitation may in general be more ubiquitous, as the action need not be contingent on body-specific properties. Thus this mechanism may not only support imitation using a different effector (e.g., a hand versus a tool or foot (Senna et al., 2014; Van Elk et al., 2011)), but also imitation between species who have different effectors (e.g., a human and a dolphin (Bauer & Johnson, 1994; Tayler & Saayman, 1973)), or species that differ vastly in size and proportion (e.g., a human and a monkey; (Kumashiro et al., 2003)). Trajectories are also a more computationally compact way of representing a movement, as they need only specify spatiotemporal parameters of the end-effector rather than information about all of the joints. Indeed, our timing data are consistent with this claim that trajectory imitation may be less computationally burdensome. Furthermore, a combined trajectory primary/intervening task resulted in the closest performance to an individual’s own baseline performance, suggesting that this combination has a lower computational demand compared to combinations of primary/intervening tasks that include posture-matching. There are also intriguing potential parallels between our results and prior research demonstrating that focusing on external, body-independent movement features (potentially akin to end-effector trajectories) is less attentionally demanding and yields greater movement automaticity compared to focusing on internal body-based movement features (e.g., (Wulf, McNevin, et al., 2001; Wulf, Shea, et al., 2001)). Finally, we note that trajectory representations seem to be readily accommodated by existing theories of motor command generation (Wong et al., 2015). Thus, while it seems that individuals are capable of imitating by matching postures or copying trajectories, future research will be necessary to further explore the conditions under which each method of imitation is favored.

It is important to point out an asymmetry in our results, in that we observed a processing cost when participants were instructed to imitate trajectories but not when instructed to imitate postures. Interestingly, this partial dissociation is consistent with another action observation study that demonstrated interference from an intervening task only in one condition (Vicary et al., 2014). We speculate that this asymmetry could have arisen for two potential reasons. The first is that there is an inherent asymmetry between the control of postures and the control of trajectories: in trajectory matching, multiple postures can be associated with a given trajectory, but in posture matching, only one trajectory can be associated with a set of postures. In other words, when imitating end-effector trajectories, due to redundant degrees of freedom at the joints it is possible to produce the same trajectory using different arm postures. Hence an intervening task that biases the use of different joint configurations is more likely to influence exactly how the arm is configured (analogous to how observed kinematics unintentionally bias one’s movement (Hardwick & Edwards, 2012; Jax & Rosenbaum, 2007)), which in turn modulates imitation accuracy. That is, the joint positions required to complete the posture matching intervening task could carry over into the trajectory primary task, disrupting performance. In contrast, moving one’s arm to achieve specific postures highly constrains the number of possible trajectories of the end-effector (in the extreme case, if one specifies the limb posture at all times during a movement, only one resulting trajectory of the end-effector is possible). As such, when the primary task was posture matching, there was a limit to how much a trajectory intervening task could bias behavior since the trajectory produced in the primary imitation task was inherently constrained by the intended postures. Hence even if trajectories are not explicitly planned when the goal is to match postures, it is much more challenging to observe a processing cost as long as nothing disrupts the representation of the desired postures themselves. This inherent asymmetry in how movements are controlled may be reflected in our findings.

A second (non-mutually-exclusive) potential explanation for only observing a processing cost when participants were instructed to imitate trajectories is that the posture-matching primary and intervening tasks were much more similar compared to the trajectory primary and intervening tasks. Specifically, both the posture matching primary and intervening task presented images of an actor to be imitated (i.e., they were both clearly imitation tasks), while the trajectory intervening task involved only viewing and tracing abstract shapes with no body information present. Thus in some sense, the trajectory intervening task could be considered more of a general motor planning task than an imitation task, compared to the posture matching intervening task. Tasks that overlap partially in terms of the underlying computations invoked tend to be most disruptive compared to tasks that overlap fully or not at all (Fournier & Richardson, 2020); thus if the trajectory intervening task invoked a set of computations that had no overlap with those required during imitation more broadly, it could explain why the trajectory intervening task was potentially less disruptive of imitation performance relative to the other conditions. Thus, there may be an inherent asymmetry in the similarity between primary and intervening tasks: one type of pairing could have induced a greater processing cost than the other pairing.

As we highlight above, the effectiveness or tendency to imitate via posture or trajectory matching is likely contingent on the task. In our paradigm we attempted to influence the type of imitation used by participants in each condition using task instructions, but it is interesting to consider what type of imitation people would choose to use by default depending on the particular movement or context (and whether that preference changes across repeated exposure or practice). As our primary imitation task involved a delay (requiring the participant to hold the movement-to-be-imitated in working memory while performing an intervening task), the imitation processes invoked in our task were likely more explicit. Additionally, our task was not a multitasking experiment *per se*, as participants were not required to perform multiple motor tasks at once. Rather, our task was a sequential multitasking paradigm (Brumby et al., 2018), where participants were required to both remember a movement while performing a different movement, then switch back to complete the imitation. It would be interesting in future work to consider how the imitation mechanism is selected during “online” imitation (when the model and actor are acting at the same time) or automatic imitation (when copying is not necessarily deliberate), and whether people can and do pay attention to different components of the movement in those situations. Moreover, despite the clear working memory demands at play, we were somewhat surprised to find no relationships between our performance measures and spatial working memory. It is possible that the spatial working memory task we used (Corsi) was not as sensitive to measuring the ability to imitate postures as a more complex sequencing or proprioception task would have been. Future research is needed to characterize individual differences in performance on imitation tasks using a variety of measures.

This initial evidence hinting at the ability to employ two distinct methods of imitation is exciting because, if borne out in future experiments, it suggests that it may be possible to modulate imitation impairments in individuals with apraxia by manipulating the focus of the task. In particular, an individual who is differentially impaired at using one of these two planning mechanisms could potentially be taught to strategically utilize the other, relatively intact planning mechanism in order to achieve the desired movement outcome (although some tasks may be more amenable to this than others). This is particularly intriguing as there is evidence that these two mechanisms may be supported by different brain regions (Buxbaum et al., 2000, 2014; Goldenberg, 2009; Wong et al., 2019), such that lesions (e.g., as a result of stroke) could differentially spare one mechanism over the other.

## Acknowledgments

The authors would like to thank Harrison Stoll for advice on the statistical approach. This research was supported by grant AES 19-02 from the Albert Einstein Society of the Einstein Healthcare Network, NIH grant R01 NS099061, NIH grant R01 NS115862, and NIH postdoctoral training fellowship 5T32HD071844.

## Declarations of interest

None.

## Footnotes

1 Segment-count errors comprised 3.33% of the data. Neither Instruction (χ^2^(3)=.97, *p*=0.8) nor the Instruction*Intervening Task interaction (χ^2^(2)=0.78, *p*=0.7) had a significant effect on segment-count error trials according to a binomial generalized linear model, but Intervening Task explained significant variance in the data (χ^2^(4)=15.50, *p*=0.004). Both Posture Matching (*z* = −3.26, *p*=0.003) and Trajectory (*z* = −3.18, *p*=0.004) ITs increased the likelihood of a segment-count error compared to the Neutral IT, but there was no difference between Posture Matching and Trajectory ITs (*z* = 0.08, *p*=0.9).

2 We ran the same analysis with the signed data, to assess whether there were systematic patterns of under- or overshooting the timing of the movement. Means are reported in Table 1. We used a linear mixed effects model, as the signed timing data were normally distributed. We observed the same patterns of results as with the unsigned data. There was a significant effect of Direction (χ^2^(1) = 95.06, *p*<0.001) but no effect of Order (χ^2^(1)=0.907, *p*=0.3).

There was a significant effect of intervening task (χ^2^(4)=30.06, *p*<0.001) but no significant interaction (χ^2^(2)=2.583, *p*=0.3) and no significant effect of Instruction (χ^2^(3)=2.767, *p*=0.4).

## Notes

### Competing Interest Statement

The authors have declared no competing interest.

### Summary of Updates

We have shortened the manuscript and strengthened the theoretical arguments. We have updated some of the figures, and cleaned up the language throughout. We have removed the secondary between-subjects consistency analysis for clarity and flow of the paper.

## References

Bates, D., Maechler, M., Bolker, B., Walker, S., Christensen, R. H., Singmann, H., & Bolker, M. (2015). Package ‘lme4’. Convergence, 12(1), 2.

Bauer, G. B., & Johnson, C. M. (1994). Trained motor imitation by bottlenose dolphins (Tursiops truncatus). Perceptual and Motor Skills, 79(3), 1307–1315.

Bekkering, H., Wohlschlager, A., & Gattis, M. (2000). Imitation of Gestures in Children is Goal-directed. The Quarterly Journal of Experimental Psychology Section A, 53(1), 153–164. https://doi.org/10.1080/713755872

Bläsing, B., Calvo-Merino, B., Cross, E. S., Jola, C., Honisch, J., & Stevens, C. J. (2012). Neurocognitive control in dance perception and performance. Acta Psychologica, 139(2), 300–308.

Bläsing, B., Puttke, M., & Schack, T. (2018). The neurocognition of dance: Mind, movement and motor skills. Routledge.

Brumby, D. P., Janssen, C. P., Kujala, T., & Salvucci, D. D. (2018). Computational models of user multitasking. Computational Interaction Design, 341–362.

Buxbaum, L. J., Giovannetti, T., & Libon, D. (2000). The Role of the Dynamic Body Schema in Praxis: Evidence from Primary Progressive Apraxia. Brain and Cognition, 44(2), 166–191. https://doi.org/10.1006/brcg.2000.1227

Buxbaum, L. J., Shapiro, A. D., & Coslett, H. B. (2014). Critical brain regions for tool-related and imitative actions: A componential analysis. Brain, 137(7), 1971–1985.

Cattaneo, L., & Rizzolatti, G. (2009). The Mirror Neuron System. Archives of Neurology, 66(5). https://doi.org/10.1001/archneurol.2009.41

Cellier, J.-M., & Eyrolle, H. (1992). Interference between switched tasks. Ergonomics, 35(1), 25–36.

Cluff, T., & Scott, S. H. (2015). Apparent and actual trajectory control depend on the behavioral context in upper limb motor tasks. Journal of Neuroscience, 35(36), 12465–12476.

Elsinger, C. L., & Rosenbaum, D. A. (2003). End posture selection in manual positioning: Evidence for feedforward modeling based on a movement choice method. Experimental Brain Research, 152(4), 499–509.

Fournier, L. R., & Richardson, B. P. (2020). Partial Repetition Between Action Plans Delays Responses to Ideomotor Compatible Stimuli [Data set]. https://research.libraries.wsu.edu:8443/xmlui/handle/2376/17908

Franz, E. A., Ford, S., & Werner, S. (2007). Brain and cognitive processes of imitation in bimanual situations: Making inferences about mirror neuron systems. Brain Research, 1145, 138–149. https://doi.org/10.1016/j.brainres.2007.01.136

Galvez-Pol, A., Forster, B., & Calvo-Merino, B. (2020). Beyond action observation: Neurobehavioral mechanisms of memory for visually perceived bodies and actions. Neuroscience & Biobehavioral Reviews.

Giese, M. A., & Poggio, T. (2003). Neural mechanisms for the recognition of biological movements. Nature Reviews Neuroscience, 4(3), 179–192.

Goldenberg, G. (1995). Imitating gestures and manipulating a mannikin—The representation of the human body in ideomotor apraxia. Neuropsychologia, 33(1), 63–72.

Goldenberg, G. (2009). Apraxia and the parietal lobes. Neuropsychologia, 47(6), 1449–1459.

Goldenberg, G. (2013). Apraxia: The Cognitive side of motor control. OUP Oxford.

Goodall, C. (1991). Procrustes Methods in the Statistical Analysis of Shape. Journal of the Royal Statistical Society: Series B (Methodological), 53(2), 285–321. https://doi.org/10.1111/j.2517-6161.1991.tb01825.x

Gower, J. C. (1975). Generalized procrustes analysis. Psychometrika, 40(1), 33–51.

Hardwick, R. M., & Edwards, M. G. (2012). Motor interference and facilitation arising from observed movement kinematics. Quarterly Journal of Experimental Psychology, 65(5), 840–847. https://doi.org/10.1080/17470218.2012.672995

Heyes, C. (2021). Imitation. Current Biology, 31(5), R228–R232. https://doi.org/10.1016/j.cub.2020.11.071

Howard, C. M., Smith, L. L., Coslett, H. B., & Buxbaum, L. J. (2019). The role of conflict, feedback, and action comprehension in monitoring of action errors: Evidence for internal and external routes. Cortex, 115, 184–200.

Isaacs, M. W., Buxbaum, L. J., & Wong, A. L. (2021). Proprioception-based movement goals support imitation and are disrupted in apraxia. BioRxiv.

Jax, S. A., Buxbaum, L. J., & Moll, A. D. (2006). Deficits in movement planning and intrinsic coordinate control in ideomotor apraxia. Journal of Cognitive Neuroscience, 18(12), 2063–2076.

Jax, S. A., & Rosenbaum, D. A. (2007). Hand path priming in manual obstacle avoidance: Evidence that the dorsal stream does not only control visually guided actions in real time. Journal of Experimental Psychology: Human Perception and Performance, 33(2), 425.

Kantak, S. S., Tessa, J., & William, M. (2020). Differential effects of internal versus external focus of instruction on action planning and performance in patients with right and left hemispheric stroke. Human Movement Science, 72, 102654.

Kumashiro, M., Ishibashi, H., Uchiyama, Y., Itakura, S., Murata, A., & Iriki, A. (2003). Natural imitation induced by joint attention in Japanese monkeys. International Journal of Psychophysiology, 50(1–2), 81–99.

Lenth, R. (2019). emmeans: Estimated marginal means, aka least-squares means. R package version 1.4. 3.01.

Matarić, M. J., & Pomplun, M. (1998). Fixation behavior in observation and imitation of human movement. Cognitive Brain Research, 7(2), 191–202.

Molenberghs, P., Cunnington, R., & Mattingley, J. B. (2009). Is the mirror neuron system involved in imitation? A short review and meta-analysis. Neuroscience & Biobehavioral Reviews, 33(7), 975–980. https://doi.org/10.1016/j.neubiorev.2009.03.010

Orban de Xivry, J.-J., & Lefèvre, P. (2016). A switching cost for motor planning. Journal of Neurophysiology, 116(6), 2857–2868. https://doi.org/10.1152/jn.00319.2016

Parsons, L. M. (1994). Temporal and kinematic properties of motor behavior reflected in mentally simulated action. Journal of Experimental Psychology: Human Perception and Performance, 20(4), 709.

Ramsay, J. R., & Riddoch, M. J. (2001). Position-matching in the upper limb: Professional ballet dancers perform with outstanding accuracy. Clinical Rehabilitation, 15(3), 324–330.

Reed, C. L., & Farah, M. J. (1995). The psychological reality of the body schema: A test with normal participants. Journal of Experimental Psychology: Human Perception and Performance, 21(2), 334.

Rohlf, F. J., & Slice, D. (1990). Extensions of the Procrustes Method for the Optimal Superimposition of Landmarks. Systematic Biology, 39(1), 40–59. https://doi.org/10.2307/2992207

Rosenbaum, D. A., Meulenbroek, R. J., Vaughan, J., & Jansen, C. (2001). Posture-based motion planning: Applications to grasping. Psychological Review, 108(4), 709.

Salvucci, D. D., Taatgen, N. A., & Borst, J. P. (2009). Toward a unified theory of the multitasking continuum: From concurrent performance to task switching, interruption, and resumption. Proceedings of the SIGCHI Conference on Human Factors in Computing Systems, 1819–1828. https://doi.org/10.1145/1518701.1518981

Schütz, C., & Schack, T. (2020). Working memory load does not affect sequential motor planning. Acta Psychologica, 208, 103091. https://doi.org/10.1016/j.actpsy.2020.103091

Schwoebel, J., Buxbaum, L. J., & Branch Coslett, H. (2004). Representations of the human body in the production and imitation of complex movements. Cognitive Neuropsychology, 21(2–4), 285–298.

Scott, S. H. (2012). The computational and neural basis of voluntary motor control and planning. Trends in Cognitive Sciences, 16(11), 541–549.

Senna, I., Bolognini, N., & Maravita, A. (2014). Grasping with the foot: Goal and motor expertise in action observation. Human Brain Mapping, 35(4), 1750–1760.

Sunderland, A., & Sluman, S.-M. (2000). Ideomotor apraxia, visuomotor control and the explicit representation of posture. Neuropsychologia, 38(7), 923–934.

Tayler, C. K., & Saayman, G. S. (1973). Imitative Behaviour By Indian Ocean Bottlenose Dolphins (t uRsiops Aduncus) in Captivity. Behaviour, 44(3–4), 286–298. https://doi.org/10.1163/156853973X00436

Team, R. C. (2013). R: A language and environment for statistical computing.

Thorndike, E. L. (1898). Animal Intelligence: An Experimental Study of the Associative Processes in Animals. Macmillan.

Tomasello, M. (1996). Do apes ape. Social Learning in Animals: The Roots of Culture, 319–346.

Van Elk, M., Van Schie, H. T., & Bekkering, H. (2011). Imitation of hand and tool actions is effector-independent. Experimental Brain Research, 214(4), 539.

Vicary, S. A., Robbins, R. A., Calvo-Merino, B., & Stevens, C. J. (2014). Recognition of dance-like actions: Memory for static posture or dynamic movement? Memory & Cognition, 42(5), 755–767.

Wong, A. L., Haith, A. M., & Krakauer, J. W. (2015). Motor planning. The Neuroscientist, 21(4), 385–398.

Wong, A. L., Jax, S. A., Smith, L. L., Buxbaum, L. J., & Krakauer, J. W. (2019). Movement imitation via an abstract trajectory representation in dorsal premotor cortex. Journal of Neuroscience, 39(17), 3320–3331.

Wulf, G. (2007). Attentional focus and motor learning: A review of 10 years of research (Target article). E-Journal Bewegung Und Training, 1, 1611.

Wulf, Gabriele. (2013). Attentional focus and motor learning: A review of 15 years. International Review of Sport and Exercise Psychology, 6(1), 77–104.

Wulf, Gabriele, Höß, M., & Prinz, W. (1998). Instructions for motor learning: Differential effects of internal versus external focus of attention. Journal of Motor Behavior, 30(2), 169–179.

Wulf, Gabriele, McNevin, N., & Shea, C. H. (2001). The automaticity of complex motor skill learning as a function of attentional focus. The Quarterly Journal of Experimental Psychology Section A, 54(4), 1143–1154.

Wulf, Gabriele, & Prinz, W. (2001). Directing attention to movement effects enhances learning: A review. Psychonomic Bulletin & Review, 8(4), 648–660.

Wulf, Gabriele, Shea, C., & Park, J.-H. (2001). Attention and motor performance: Preferences for and advantages of an external focus. Research Quarterly for Exercise and Sport, 72(4), 335–344.

Zentall, T. R. (2006). Imitation: Definitions, evidence, and mechanisms. Animal Cognition, 9(4), 335–353. https://doi.org/10.1007/s10071-006-0039-2

